# Automated sequential chromogenic IHC double staining with two HRP substrates

**DOI:** 10.1101/276501

**Authors:** Kenneth H Petersen, Jesper Lohse, Lasse Ramsgaard

## Abstract

Automated IHC double staining using DAB and HRP Magenta is illustrated utilizing a new acid block with sulfuric acid to prevent cross-reactivity. Residual cross-reactivity in double staining is determined to arise from chromogenic-bound antibodies and amplification system during the first part of the double staining.

## Introduction

A concern with many diagnostic procedures today is to get enough information from limited sample sizes to reach an informed diagnostic decision. In pathology, using immunohistochemistry (IHC), this is particularly important when the biopsy is small and only a few different tests can be run on the available tissue. The localization of different antigens in relation to each other can in some cases also be important for a diagnosis. In these cases, staining the same tissue section for two antigens is a useful diagnostic tool.

To reduce the possibility of cross-staining between the two targets, antibodies derived from different species, usually rabbit and mouse, are commonly used (van der Loos 1993). Additionally, the visualization of the antigens is performed using different enzymes in the visualization steps. Typically, brown and red visualization is performed using the enzymes horseradish peroxidase (HRP) and alkaline phosphatase (AP) and the substrates diaminobenzidine (DAB) and Fast Red, respectively (Malik 1982). No quenching of enzymatic activity is required between the two stains as the two enzymes do not function with each other’s substrate. Both colors contrast well to the commonly used blue hematoxylin nuclear stain.

However, there are drawbacks to the AP substrate system. First, Fast Red stains rapidly dissolve in ethanol and most organic mounting media. Second, the Fast Red diazonium salt can react with a range of other substances present in the tissue section. This includes the DAB stain which becomes darker after incubation with Fast Red. Third, Fast Red chromogen must be used within 30 minutes of mixing. For automated staining systems, this requires onboard mixing of the Fast Red reagents or alternatively the staining is paused until the user can supply a freshly mixed Fast Red solution to the system.

An alternative red chromogen, 3-amino-9-ethylcarbazole (AEC) exists for the HRP enzyme system, but this is not compatible with other peroxidase substrates due to poor color contrast between reddish-brown AEC and brown DAB (Nemes 1987).

Blue HRP substrates have previously been described. While they contrast well to DAB they are not optimal when hematoxylin is used as a counterstain (Petersen 2009).

Recently a new magenta-colored HRP chromogen with a sensitivity comparable to DAB has been described (Lohse 2016). This paper reports our findings with fully automated DAB/Magenta chromogen IHC double stains in combination with hematoxylin nuclear counterstain.

## Results and discussion

When performing double staining, it is important that any cross-reactivity between reagents is eliminated. In this study, we used two chromogens that are both substrates for the HRP enzyme. This makes it important to efficiently remove all enzyme activity from the first stain before the second stain. Peroxidase blocking is normally used to quench any endogenous peroxidase activity in the tissue (Li 1987). However, this does not remove all HRP activity when the tissue contains additional peroxidase activity from the visualization conjugates as seen in Figure 1B, thus additional removal of peroxidase activity is needed. Sequential double staining using two HRP enzymes has previously been described using an acidic block between the two stains (Nakane 1968). An acid block works by dissociating the antibodies from their targets so they can be washed away.

**Figure 1.**
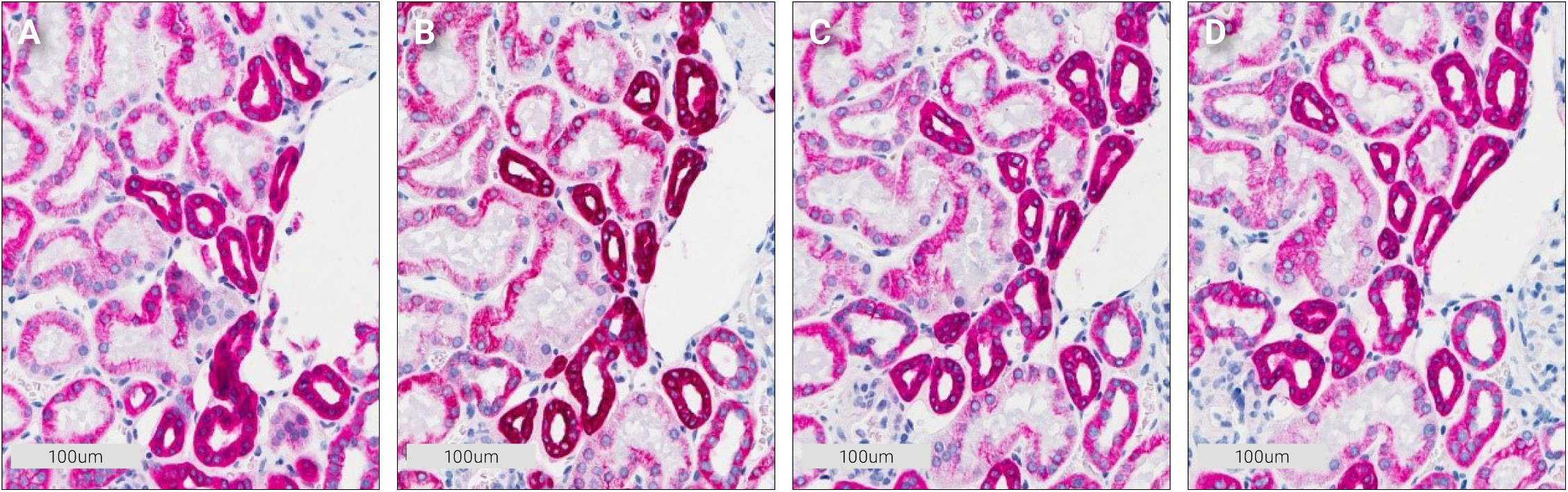
Effect of different blocking washes on cytokeratin Pan staining in kidney tissue. **A** No wash, No DAB. **B** Peroxidase block then DAB. C 50 mM H_2_SO_4_ then DAB. D 300 mM H_2_SO_4_ then DAB.

### Sulfuric acid treatment

The efficiency of using an acidic block was tested using sulfuric acid. Starting from 300 mM sulfuric acid a two-fold dilution series down to 20 mM sulfuric acid was made and tested with four different antibodies. The acid block step was set to 3 minutes, which is the minimum time on the automated Dako Omnis staining system for a block or incubation step. The test was set up as a single chromogenic stain with HRP Magenta chromogen followed by a 3-minute acid block, wash buffer and then a DAB chromogen incubation. No DAB staining was visible on any of the slides, the acid block removed all detectable enzyme activity and the morphology of the tissue was not visibly impacted (Figure 1C+D).

Next, we investigated if either of the two chromogens from a completed staining was affected by a sulfuric acid block. A Ki-67 stain of nuclei in tonsil and colon tissue was used for this experiment. Ki-67-stained nuclei can be counted using an image analysis algorithm. Slides stained with either DAB or HRP Magenta were subjected to a subsequent block with sulfuric acid, in concentrations ranging from 0 to 400 mM. The percentage of Ki-67 positive nuclei in tonsil and colon was counted for each concentration of sulfuric acid. Figure 2 shows that the tested concentrations have no effect on the staining of the slides, the chromogen already present on the slides is not removed or faded by the acid block. We conclude that sulfuric acid in the range 50-400 mM effectively removes residual HRP activity, without negatively impacting the intensity of a first DAB or HRP Magenta stain.

**Figure 2.**
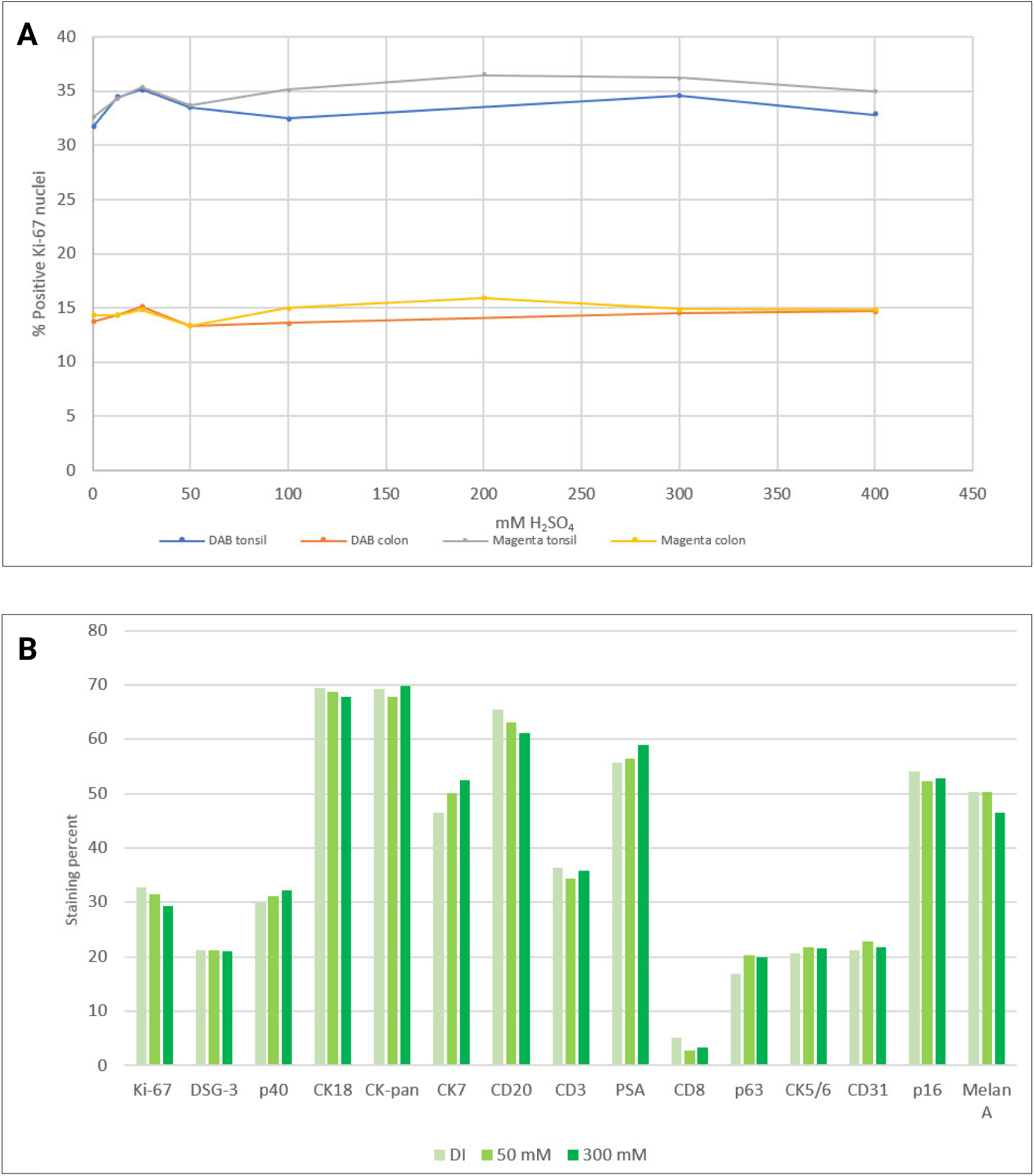
A Percent Ki-67 positive nuclei in tonsil and colon using either DAB or Magenta as chromogen. **B** Effect of sulfuric acid block before IHC staining and Magenta as chromogen reported as either percent stained nuclei or percent stained complete membranes (DI=deionized water).

Some epitopes may be sensitive to the sulfuric acid block which could affect the intensity of the second staining. To investigate this, we applied an acid step prior to application of the primary antibody and completed the staining as a single stain with either DAB or HRP Magenta chromogen. Three different treatments before primary antibody were compared: Block with deionized water, 50 mM or 300 mM sulfuric acid. A range of different antibodies were tested and the stains were evaluated using digital image analysis. As seen in Figure 2B, we observed minor changes in staining intensity which we consider to be within the uncertainty of the experiment. Here we use the stains qualitatively and the effect (if any) is therefore negligible. It is, however, important to be aware of any impact on staining intensity from either the first staining or the acid treatment if a double staining is to be used for quantification.

Moving forward with the double staining, we decided to use a belt-and-braces approach with sulfuric acid first, followed by a peroxidase block to ensure that all cross-reactivity was eliminated. The two quenching reagents are orthogonal; acid block dissociates the antibodies from their targets while peroxidase block works by “overloading” the enzymes with hydrogen peroxide they cannot get rid of as no substrate is present. This eventually quenches the enzyme activity. Sulfuric acid is placed first in the second part of the staining protocol to remove as much bound antibody as possible before any remaining antibody-enzyme conjugates still in the tissue are quenched by treatment with the peroxidase block.

### Non co-localized targets

We performed a double staining with monoclonal mouse antibodies against a nuclear (p63) and a membrane target (carcinoembryonic antigen (CEA)), respectively. The guiding principle being that a first and a second IHC stain including the subsequent hematoxylin counterstain should ideally stand out unaffected by each other since they are not co-localized. Staining using the same combination of antibodies were run both with HRP Magenta stain first then DAB and vice versa. Figure 3A-D shows these double stains.

**Figure 3.**
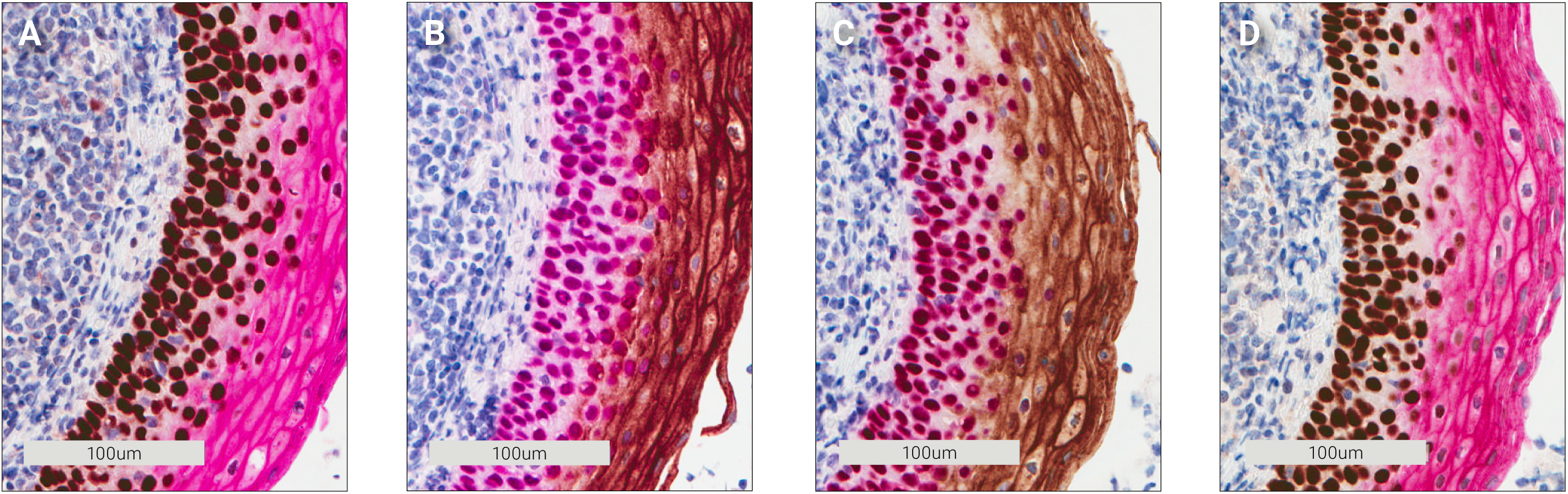
Carcinoembryonic antigen (CEA) and p63 double staining of tonsil tissue using monoclonal mouse antibodies. **A** 1st. p63/DAB 2nd. CEA/ Magenta. **B** 1st. CEA/DAB 2nd. p63/Magenta. **C** 1st. p63/Magenta 2nd. CEA/DAB. **D** 1st. CEA/Magenta 2nd. p63/DAB.

Cross-reactivity can be seen when DAB is used as the first chromogen, either as a reddish ring around the nuclei (Figure 3A) or as a change in the brown color (Figure 3B). Likewise, when the magenta color is used first, the magenta color becomes a darker red (Figure 3C+D). The purple color of the nuclei in Figure 3B is expected since it is a combination of the magenta and the blue hematoxylin colors.

The observed color spillover was not expected because our first experiments demonstrated that we had removed all peroxidase activity in the tissue. The reason for the cross-reactivity must be found by looking beyond the enzyme. We believe an explanation for this unexpected color spillover is that reagents from the first stain remain in the tissue because the chromogen has covalently bonded them to the tissue (Figure 4A-E). When the second stain is performed the reagents are recognized by remaining amplification system (Figure 4C) and remaining primary antibodies are recognized by the amplification system along with the primary antibodies captured by the remaining amplification system (Figure 4D). During the second chromogen incubation the chromogen is then precipitated (Figure 4E) and this gives rise to the color spillover.

**Figure 4.**
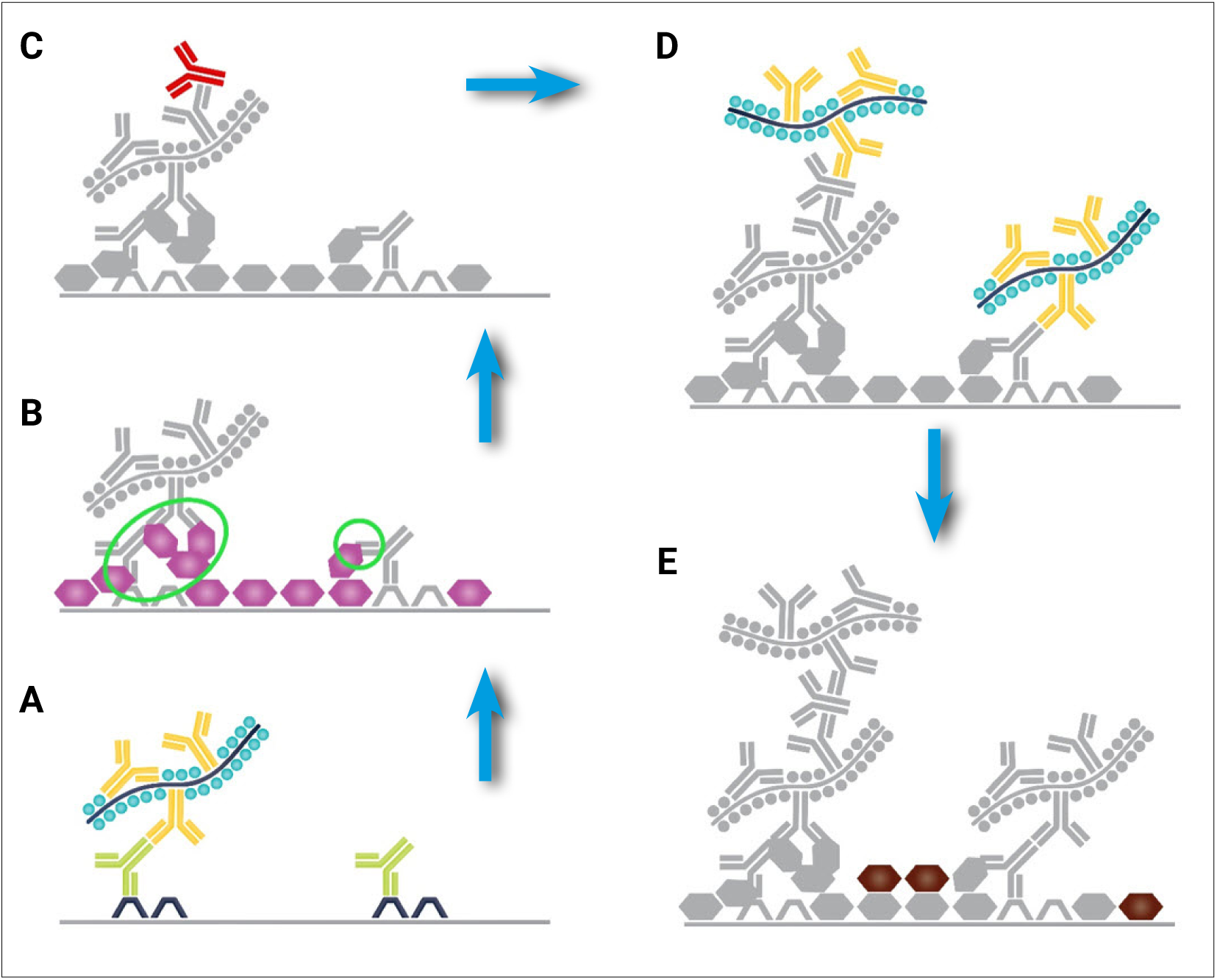
Cross-reactivity through irreversible binding of antibodies. **A** The first primary antibody recognizes the first target and is recognized by the amplification system. **B** Chromogen is precipitated and covalently binds antibodies and amplification system to the tissue, green circles. **C** The second primary antibody is captured by free antibodies of the amplification system. **D** As the second layer of amplification is applied, antibodies captured by the first amplification layer along with those covalently bound to the tissue are captured. **E** And color spillover results when the next chromogen is applied.

To test our theory, we expanded on our initial sulfuric acid and peroxidase block experiments. First a complete stain for cytokeratin (CK) Pan with HRP Magenta was done and then the slide was treated with 300 mM sulfuric acid and peroxidase block to remove all remaining peroxidase activity. Next, we incubated with the amplification system once more before a final DAB incubation. As seen in Figure 5A-B, the magenta color becomes visibly tainted by DAB (it is darker). We believe this supports the theory.

**Figure 5.**
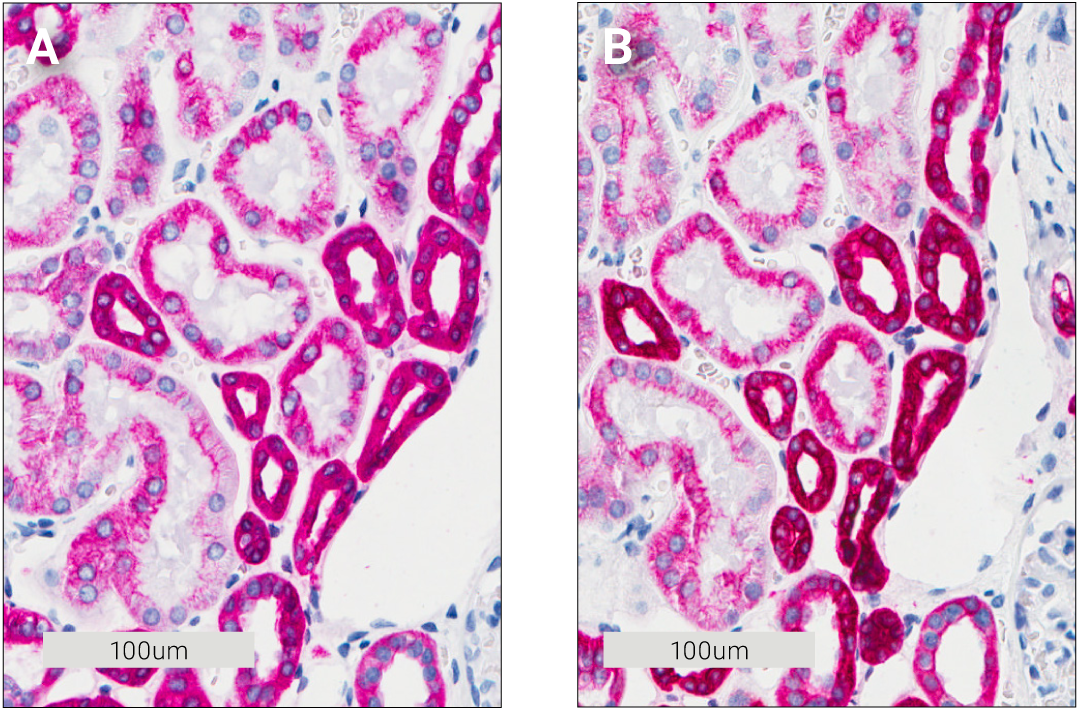
Demonstration of residual covalently bound antibodies in kidney tissue after sulfuric acid and peroxidase block treatment by incubation with DAB after block. **A** 300 mM sulfuric acid block. **B** 300 mM sulfuric acid, peroxidase block then amplification.

The double staining of p63 and CEA was repeated; this time we used a rabbit antibody against CEA while the p63 antibody was the same mouse antibody as in the initial experiment. As seen in Figure 6A-D the colors of the double stainings became untainted, regardless of which chromogen was used as the first chromogen (compare Figure 3B with 6B and Figure 3D with 6D).

**Figure 6.**
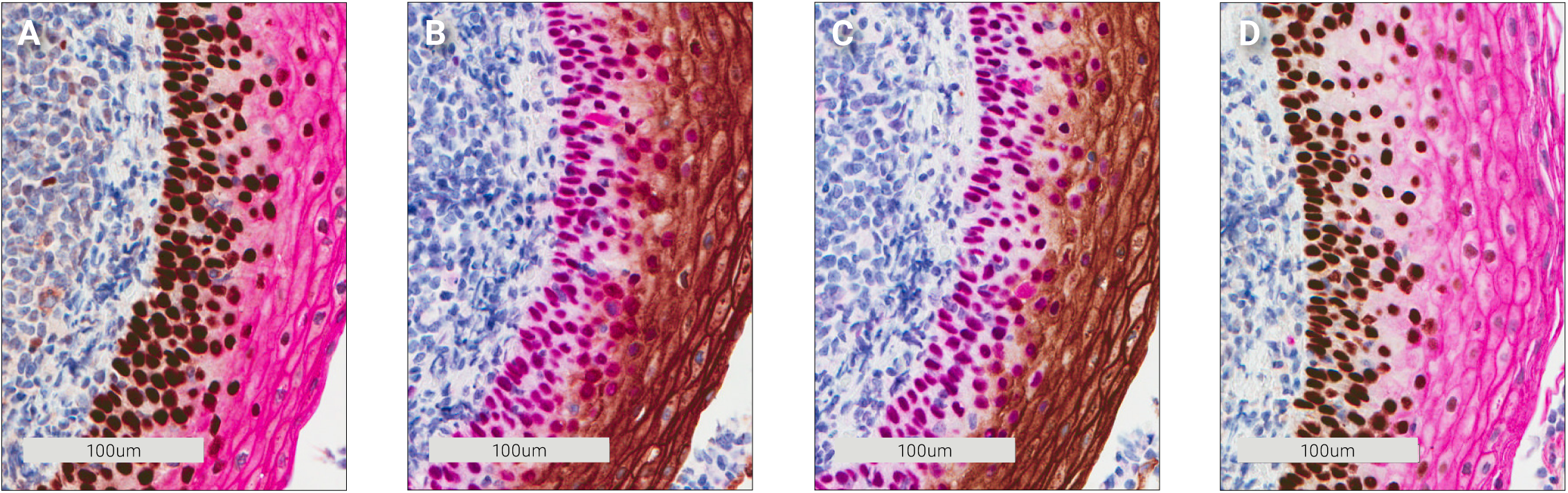
Rabbit polyclonal CEA and mouse monoclonal p63 antibody double staining of tonsil tissue. **A** 1st. p63/DAB 2nd. CEA/Magenta. **B** 1st. CEA/ DAB 2nd. p63/Magenta. **C** 1st. p63/Magenta 2nd. CEA/DAB. **D** 1st. CEA/Magenta 2nd. p63 DAB.

### Co-localized targets

Our next step was to investigate how a double stain would look if the targets were co-localized. For this experiment, we used cytokeratin (CK) Pan and cytokeratin 18 antibodies. CK 18 is not recognized by the CK Pan antibody, but is co- localized with CK Pan in prostate glands. Figure 7A-D shows the result from the cytokeratin double stainings. The CK Pan antibody stains a slightly larger area of the prostate epithelium than CK 18. This gives a rim of either HRP Magenta stained CK Pan (Figure 7A) or DAB-stained CK Pan around the CK 18 stain (Figure 7C). The rim is only visible when CK 18 is used as the first antibody in the double staining. If the larger CK Pan area is stained first, a shielding of the co-localized CK 18 target then comes into effect. This markedly lowers the staining intensity of CK 18. In the case where DAB is used together with CK Pan, the co-localization of the second target is not easily seen (Figure 7B). The transparency of the magenta color makes it comparably easier to see that the targets are co-localized. The HRP Magenta/DAB combination is not ideal for co-localized targets, but using the more transparent HRP Magenta as the first chromogen increases the likelihood that co-localized targets are detected due to tainting of the magenta color. When DAB was used as the first chromogen only the expected areas for each antibody were stained and the colors are not visibly tainted. We ascribe the observed difference to the shielding effect that DABhas on epitopes that are stained with the DAB chromogen. The DAB precipitate can be so dense that access to epitopes in the tissue by subsequent antibodies is almost impossible. This combined with the ability of the brown color to dominate other colors keeps the DAB stain brown (Figure 7A) even after the second staining is complete. The same is true in Figure 7B where the DAB stain keeps its brown appearance but is lighter due to some HRP Magenta precipitation. We ascribe this to different expression levels of cytokeratin, which affects how dense the DAB precipitate becomes.

**Figure 7.**
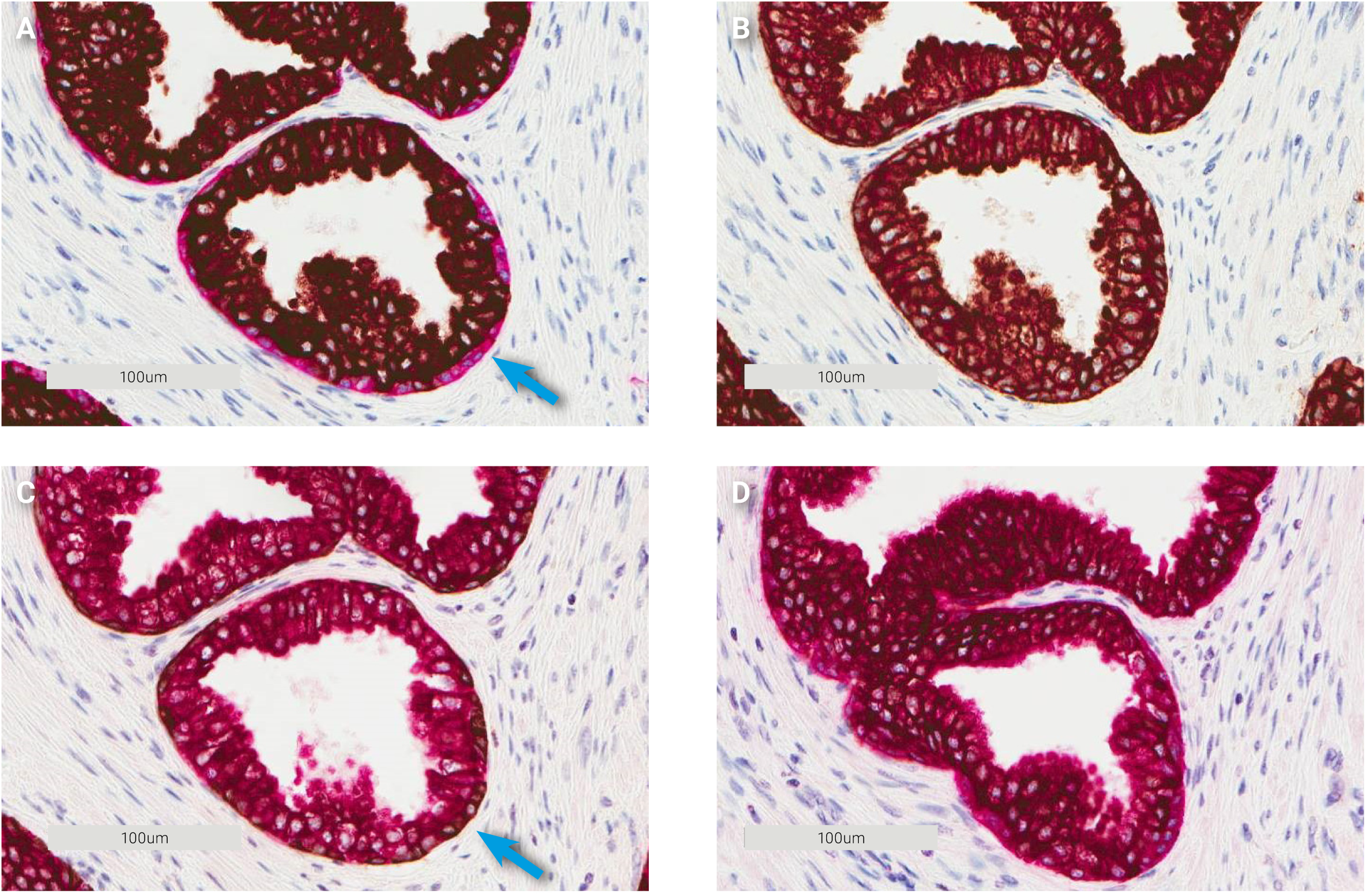
CK Pan/ CK 18 double staining of prostate tissue on Dako Omnis. **A** 1st. CK 18/DAB 2nd. CK Pan/Magenta. **B** 1st. CK Pan/DAB 2nd. CK 18/ Magenta. **C** 1st. CK 18/Magenta 2nd. CK Pan/DAB. D 1st. CK Pan/Magenta 2nd. CK 18/DAB.

For the HRP Magenta chromogen the situation is the same, a high shielding effect is observed when CK 18 is stained first and less shielding when CK Pan is stained first.

## Conclusion

We have demonstrated double staining using two HRP substrates as chromogens. The HRP activity from the first staining can be completely removed by a sulfuric acid block step. Using the HRP Magenta chromogen in combination with DAB makes fully automated protocols for double staining possible.

The best double staining in terms of no cross-reactivity is still made with antibodies from two different species. However, in cases of densely packed epitopes using DAB as the first chromogen, the spillover is practically hidden by the intense DAB stain. This makes it possible to do a double staining with two antibodies from the same species. Using antibodies from different species the sequence of the chromogens is not important. Both DAB and the HRP Magenta chromogen can be used as a first stain. We have explained why cross-reactivity remains an issue when using antibodies from the same species.

In our experience the use of DAB as the first chromogen and HRP Magenta as the second chromogen have given consistently good results with minimal detectable cross-reactivity and this is now our standard approach when setting up double staining protocols.

## Materials and Methods

All reagents were purchased from Sigma Aldrich and Agilent Technologies. Antibodies were purchased from Agilent Technologies and used according to the manufacturer’s specifications. Desmoglein-3, p40 and p16 were from Abcam, Biocare and Santa Cruz Biotechnology. Automated staining was performed on either a Dako Omnis system or an Autostainer Link 48 system. On the Dako Omnis system the “IHC Double Stain Template” was used. For the Autostainer Link 48, a protocol was defined and it can be found in the supplementary material.

Images were captured on an Aperio Scanscope. Areas of interest were loosely circled and analyzed using the membrane v9 or the nuclear v9 algorithm adjusted to the color of the chromogen.

### Specimens

All tissue was obtained from Department of Pathology, Odense University Hospital, Region South, Denmark, fixed in formaldehyde for 24 hours prior to embedding in paraffin. All specimens were completely anonymized prior to receipt at Agilent. According to the Danish law on the Research Ethics Committee System and handling of biomedical research projects and communication between Agilent and the Danish Committee on Biomedical Research Ethics and the Regional Ethics Committee (IRB), the tests performed at Agilent on anonymous residual tissue are not subject to an approval by the IRB system because such studies are considered quality control projects. Therefore, no IRB approval for this work has been obtained.

## Acknowledgements

The Technical assistance of Anne Bruun, Kristine Faerk, Maria Kristensen, Anne Maarbjerg and Marianne Marcussen is gratefully acknowledged.

